# *Death & Chemotaxis*: Bacterial chemotaxis enables collective escape from phage predation

**DOI:** 10.64898/2026.01.28.702367

**Authors:** Victoria G. Muir, Alejandro Martínez-Calvo, Ned S. Wingreen, Sujit S. Datta

## Abstract

Bacteriophages (“phage”) are viruses that prey on bacteria in diverse environments, from biological tissues to soils. In many of these environments, bacterial hosts are constantly migrating, yet how bacterial migration is influenced by phage predation remains poorly understood. Using transparent granular hydrogels that mimic natural habitats, we directly visualize populations of motile Escherichia *coli* encountering lytic T4 phage. Unexpectedly, we find that even in phage-rich environments, bacteria successfully form chemotactic fronts that enable them to migrate over large distances without needing to develop phage resistance. Higher phage concentrations delay front formation but not steady-state front speed or shape. By combining our experiments with biophysical modeling, we demonstrate that this phenomenon arises from the ability of cells to collectively outrun trailing phage bursts—as quantified by a dimensionless “escape parameter” comparing chemotactic and predation rates. This work thus reveals and provides mechanistic insight into the role of cell motility in shaping phage-bacteria interactions in spatially-extended environments.

---

Phage are the most abundant biological entities on Earth, shaping bacterial behavior in diverse settings ranging from the human body to terrestrial and marine ecosystems [1–7]. Traditional laboratory studies examine phage-bacteria interactions in well-mixed liquid cultures, which eliminate spatial structure, or on agar plates, where the focus is typically on infection efficiency and resistance evolution [8–10]. However, in nature, bacteria migrate collectively over large distances via chemotaxis [11, 12], navigating chemical gradients to find nutrients, colonize hosts, cause infections, and regulate biogeochemical processes [12–17]—all while encountering phage predators [7]. The ubiquity of this process thus presents a puzzle: How do bacteria successfully traverse these environments where phage are abundant—even outnumbering bacterial cells by up to an order of magnitude [6, 18–21]—without being decimated?

Resolving this puzzle requires understanding the interplay between bacterial collective migration and phage predation, which remains poorly understood. Recent work has begun to address this gap in knowledge by studying how phage can “hitchhike” on migrating bacteria to co-propagate through space and form characteristic spatial infection patterns with their motile hosts [22–24] and influence co-evolutionary dynamics [25]. Yet a fundamental question remains: Can bacteria traverse phage-rich environments without needing to evolve resistance, and if so, what biophysical principles govern population survival?

Here, we show that they can—by outrunning phage. Using transparent granular hydrogels that mimic natural porous habitats [26, 27], we directly visualize populations of motile *E. coli* encountering lytic T4 phage. By consuming chemoattractant, the bacteria establish a self-generated gradient that drives collective chemotaxis, forming migrating fronts even at the highest phage concentrations. By integrating experiments and modeling, we show that while phage continually infect and lyse a subpopulation of cells, producing “bursts” of new phage progeny, the collective migration is fast enough for the population to outpace the spreading infection—enabling long-range migration without the need to evolve resistance. This work thus reveals an ecological role of chemotaxis beyond nutrient foraging and range expansion: it enables bacteria to escape phage predators in spatially-extended environments, establishing quantitative principles for predicting population survival versus collapse.

## Results

### Phage delay chemotactic front formation without affecting steady-state shape or speed

Our experiments explore the collective migration in glass capillaries of motile *E. coli* through granular hydrogel packings, with defined concentrations *p*_init_ of T4 phage mixed in [Fig. **1**a, *Methods*]. Each packing is made of jammed [Fig. S1], biocompatible, ∼10 μmdiameter hydrogel particles swollen in a defined rich liquid medium with glucose and *L*-serine as the primary nutrient and chemoattractant, respectively. The internal mesh size of each hydrogel particle is ≈ 40 − 100 nm [28], large enough to allow unimpeded transport of oxygen and nutrient [29–32], but smaller than both cells and phage [33]. Therefore, neither cells nor phage can penetrate the individual particles; the cells are instead forced to swim through the pores formed in the interstices between particles [26, 34], while the phage randomly diffuse through this pore space, remaining stable [Fig. S2] and lytic [Fig. S3] for at least 48 h, much longer than the experimental duration. Moreover, because the hydrogel particles are so highly swollen with liquid, the packings are transparent— enabling us to use fluorescence confocal microscopy to directly visualize the cells, which constitutively express GFP in their cytoplasm, as they move in situ [26, 34].

**Fig. 1.**
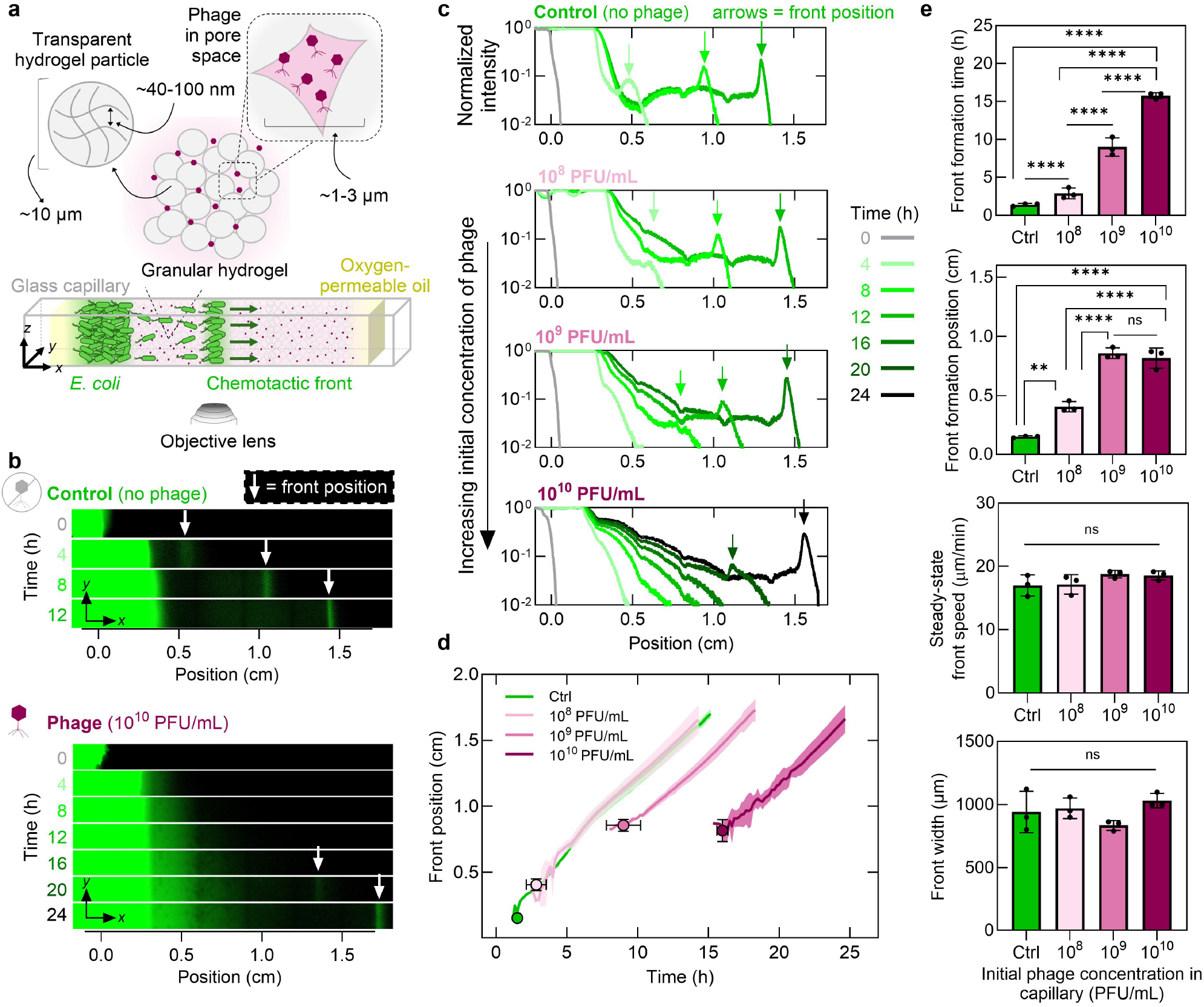
The presence of phage delays chemotactic front formation but does not alter its steady-state dynamics. **a**, Schematic of experimental setup. Fluorescent *E. coli* migrate from a dense inoculum placed at one end of a glass capillary through the pores of a granular packing of hydrogel particles. The liquid-infused particles are transparent and permeable to nutrients and small molecules but not to bacteria or phage; therefore, phage diffuse through the pore space and infect cells. We cap each end of the capillary with oxygen-permeable paraffin oil to prevent evaporation and maintain the entire system at 30°C. **b**, Confocal micrographs (depth *z*-integrated fluorescence signal) showing chemotactic front migration along the capillary for a phage-free control (top) and high phage concentration (bottom). **c**, Normalized fluorescence intensity (integrated along *y* and *z*) profiles along the capillary for control, 10^8^ PFU/mL, 10^9^ PFU/mL, and 10^10^ PFU/mL phage concentrations. Arrows indicate front location at given times. **d**, Chemotactic front position over time for various phage concentrations. Solid curves show average front positions; shaded regions indicate standard deviation. Circles denote average front position at formation time; vertical and horizontal error bars show standard deviations. **e**, Quantification of front formation time, formation position, steady-state speed, and width at various phage concentrations. Data represent mean ± standard deviation (*n* ≥ 3). One-way ANOVA: ns = not significant, \*\**p* < 0.01, \*\*\*\**p* < 0.0001.

In each experiment, we place a dense inoculum of cells (concentration *b*_init_ ∼ 10^11^ CFU/mL) at one end of a capillary (*x* ≈ 0) and track their migration to the other end through the hydrogel packing in real time *t* [Video S1]. First, we examine the phage-free case, which serves as a negative control. The cells in the inoculum continually consume the chemoattractant, establishing a local gradient that biases their motion via chemotaxis—eventually forming a localized ∼1000 μm-wide front of migrating cells after ≈ 2 h [arrows, Fig. **1**b-c (top)]. The continued consumption of chemoattractant by these cells causes the chemotactic front to continue to propagate through the capillary at a constant speed of ≈ 17 μm/min [green curve, Fig. **1**d (top)], consistent with the findings of previous studies [17, 27, 35, 36]. The fluorescence signal intensity in the migrating front increases over time [Figs. **1**b, S4], reflecting the growth of new cells that compensates for leakage from the front, sustaining steady-state migration of the front with bacterial concentration *b* ∼ 10^10^ − 10^11^ CFU/mL [37].

Next, we repeat the same experiment, but with phage uniformly dispersed through the hydrogel packings at increasing concentrations *p*_init_ = 10^8^, 10^9^, and 10^10^ PFU/mL, corresponding to an initial effective multiplicity of infection MOI_init_ ≡ *p*_init_*/b*_init_ of 10^−2^ − 10^0^. These conditions approximate those found in natural environments [6, 18–21], though unlike in well-mixed laboratory cultures, bacteria and phage are initially spatially separated here. Remarkably, even under this extreme predation pressure—which rapidly decimates the population in well-mixed culture [Fig. S3]—the bacteria form migrating chemotactic fronts in the spatially-extended system, even at the highest phage concentration [Fig. **1**b (bottom)]. The time required for each front to form and the initial position of detectable front formation both increase with phage concentration [Fig. **1**e (top two panels)]. Yet, once a chemotactic front forms, it traverses the phage-rich terrain with a steady-state concentration, speed, and width that are no different from the phage-free case [Figs. **1**b-e (lower two panels), S4]—despite the local MOI within the front being as large as ∼ 10^0^, corresponding to one phage per bacterium.

### Phage exposure enriches for bacteria with reduced phage susceptibility in chemotactic fronts

Given the strong selection pressure imposed by continuous phage predation in our experiments, one possible reason why bacteria still survive and form chemotactic fronts is that the cells evolve phage resistance [38]. To test this possibility, we isolate bacteria from chemotactic fronts after they have traversed 1 cm of the phage-laden hydrogel [Fig. **2**a]—representing hours of continuous phage exposure across multiple bacterial generations. We dilute and plate these cells on soft agar that is either phage-free or phage-rich; each resulting colony-forming unit (CFU) then originates from a single cell within the front [Fig. S5].

**Fig. 2.**
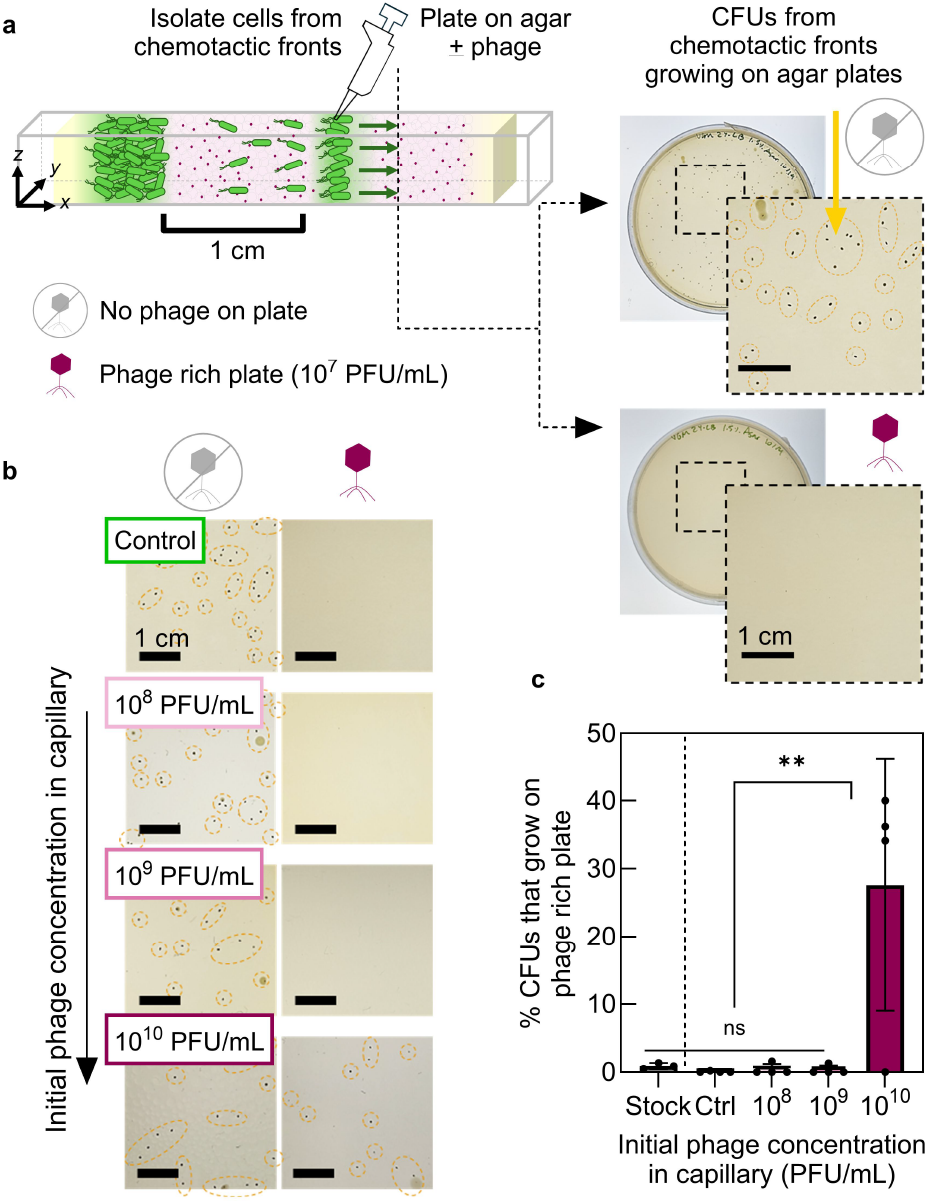
High phage exposure enriches for bacteria with reduced phage susceptibility in chemotactic fronts. **a**, Schematic of isolating cells from chemotactic fronts after ∼ 1 cm migration. The cells are collected and diluted onto soft agar plates lacking phage (top) or containing phage at 10^7^ PFU/mL (bottom). Scale bar = 1 cm. **b**, Representative images of colony-forming units (CFUs) from bacteria isolated from chemotactic fronts. Left: CFUs on phage-free plates. Right: CFUs (or absence) on phage-rich plates, depending on initial phage concentration during front formation. Scale bar = 1 cm. **c**, Percentage of bacteria from chemotactic fronts that grow in phage-rich environment, calculated by dividing CFUs on phage-rich plates by CFUs on phage-free plates. Data represent mean ± standard deviation (*n* ≥ 3). One-way ANOVA: ns = not significant, \*\**p* < 0.01.

The number of CFUs on phage-free plates remains consistent across all conditions tested [Figs. **2**b, S5], indicating that the concentration of cells in the front is the same across experiments. Importantly, none of the cells from phage-free control fronts form colonies on phage-rich plates [Fig. **2**b-c], confirming that the inoculated population lacks pre-existing resistance. Moreover, cells isolated from fronts formed at the lower phage concentrations (10^8^ and 10^9^ PFU/mL) similarly cannot form colonies on phage-rich plates [Fig. **2**b-c]—indicating that, despite traversing the phage-rich terrain in a chemotactic front, most cells remain fully susceptible to phage infection. By contrast, ≈ 30% of cells from fronts exposed to the highest phage concentration (10^10^ PFU/mL) form colonies on phage-rich plates [Fig. **2**b-c]. This finding suggests that prolonged, intense phage exposure can enrich for cells with reduced susceptibility; however, even in this extreme case, the majority of cells in the migrating front remain susceptible to phage. Thus, resistance evolution alone cannot explain how bacterial populations successfully traverse phage-rich environments via chemotaxis.

### Biophysical model reveals that collective chemotaxis enables bacteria to outrun phage bursts

If resistance does not explain bacterial survival, what does? To understand how chemotactic bacteria successfully traverse phage-rich environments, we develop a biophysical model incorporating the key features of our experimental system. All quantities are averaged along the transverse (*y*-*z*) direction, reducing the model to a one-dimensional system with space parameterized by the *x* coordinate. As summarized in Fig. **3**a, we consider a population of motile, growing bacteria migrating into an initially uniform suspension of lytic phage, which are represented by the concentration field *p*(*x, t*) and have diffusivity *D*_*p*_. Adsorption of phage then converts uninfected cells into infected cells (denoted by the subscript _inf_), which become non-motile [39] and lyse at a rate *δ*, releasing a burst of *β* new phage.

**Fig. 3.**
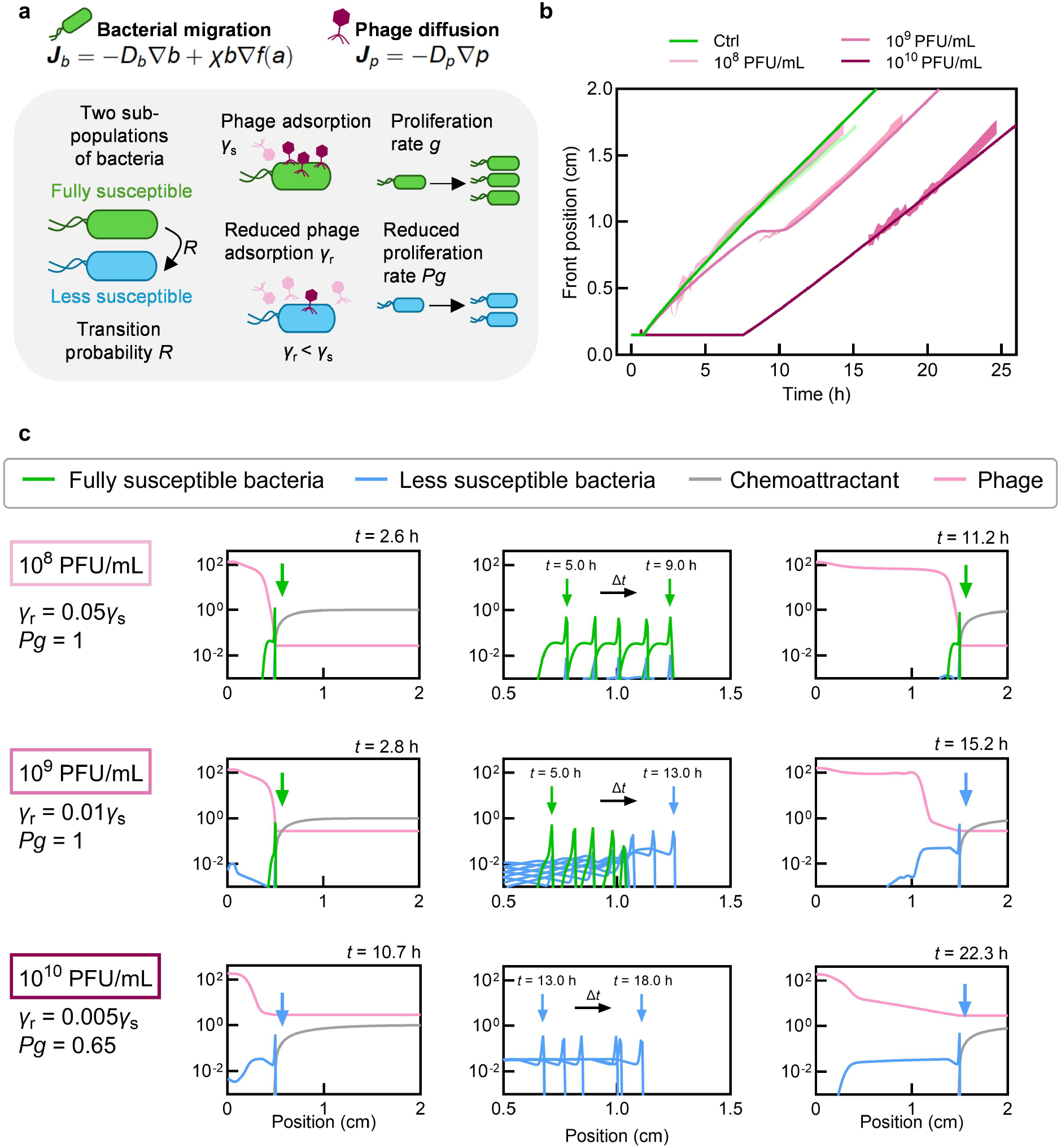
Biophysical model reproduces the essential features of bacterial collective chemotaxis in the presence of phage. **a**, Model schematic showing two swimming bacterial subpopulations: fully susceptible (proliferating at rate *g*) and less susceptible (proliferating at reduced rate *Pg*, with *P* < 1). Fully susceptible bacteria transition to less susceptible at rate *R*. Both populations consume nutrients and perform chemotaxis with identical characteristics; phage infect these populations at rates *γ*_s_ and *γ*_r_ < *γ*_s_. **b**, Bacterial chemotactic front position versus time for phage-free environment and *p*_init_ = 10^8^, 10^9^, and 10^10^ PFU/mL, from model simulations (solid curves) and experiments (shaded regions show standard deviation). **c**, Normalized concentration profiles of bacteria, chemoattractant, and phage at different times for various phage concentrations. Bacteria and chemoattractant profiles are normalized by their initial concentrations at *t* = 0, and the phage profile is normalized by the characteristic concentration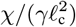, where ℓ_c_ is the chemoattractant penetration scale (see Table **1** and Methods). Initially, nutrient and chemoattractant are uniform across the domain with concentrations *c*_init_ and *s*_init_, respectively. Bacteria are uniformly distributed in a phage-free region (0 < *x* < *x*_init_) adjacent to uniform phage suspension (*x*_init_ < *x* < *L*), forming a slab-shaped colony facing the phage-rich environment as in the experiments. Here, *x*_init_ denotes the position of the bacteria–phage contact interface at *t* = 0, and *L* is the domain length. We assume phage are stable and long-lived over experimental duration, consistent with observations (Fig. S2). At low phage concentration (10^8^ PFU/mL), bacteria rapidly establish a chemoattractant gradient and form a migrating front, leaving phage behind. At intermediate concentration (10^9^ PFU/mL), phage initially decimate the susceptible population; the less susceptible subpopulation then proliferates and reestablishes the gradient before migrating. At high concentration (10^1^0 PFU/mL), this delay is further prolonged but the front eventually forms and migrates at the same steady-state speed. Arrows indicate chemotactic front position.

Bacterial populations naturally exhibit heterogeneity in susceptibility to phage, arising from purely phenotypic variation in receptor expression and phage adsorption rates [40–42]. To capture this variability in a simplified manner, we classify the bacteria into two subpopulations *b*_s_(*x, t*) and *b*_r_(*x, t*): fully susceptible cells with infection rate *γ*_s_ (99% of the initial population) and less susceptible cells with a reduced infection rate *γ*_r_ < *γ*_s_ (1%) due to reduced phage adsorption, respectively. Motivated by our experimental findings, we assume that neither subpopulation acquires mutations conferring phage resistance during migration. In addition, to capture phenotypic plasticity in phage susceptibility [43–45], we include a small transition rate *R* for susceptible cells to become less susceptible. We also incorporate a growth penalty *P* ∈ (0, 1) for less susceptible cells to describe any fitness loss as a consequence of becoming less susceptible to phage infection [46, 47].

Both bacterial subpopulations consume nutrient *c*(*x, t*), which has diffusivity *D*_*c*_ and enables the cells to grow at a maximal rate *g* modulated by the Michaelis-Menten function *f*_g_(*c*) = *c/*(*c* + *c*_M_) with characteristic concentration *c*_M_. They also consume chemoattractant *a*(*x, t*), which has diffusivity *D*_*a*_, creating a self-generated gradient that drives chemotactic migration with flux ***J***_*i*_ = −*D*_*b*_**∇***b*_*i*_ + *b*_*i*_***v***_chemo_. The first term describes unbiased random motion of cells with effective diffusivity *D*_*b*_, and the second describes their directed chemotaxis with velocity ***v***_chemo_ = *χ***∇***f*_a_(*a*). Here, *f*_a_(*a*) = ln[(*a* + *a/a*_−_)*/*(*a* + *a/a*_+_)] describes the ability of the bacteria to logarithmically sense the chemoattractant [17, 35, 36, 48, 49] with upper and lower characteristic concentrations *a*_+_ and *a*_−_, respectively, and the chemotactic coefficient *χ* describes their ability to move up the sensed gradient. We assume that both nutrient and chemoattractant are consumed at a maximal rate *k*, also following the same Michaelis-Menten kinetics as growth.

Altogether, these processes are summarized by the following dynamical equations:

Susceptible bacteria:

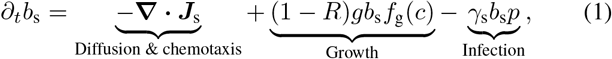

Infected susceptible bacteria:

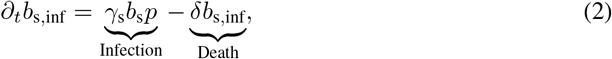

Less susceptible bacteria:

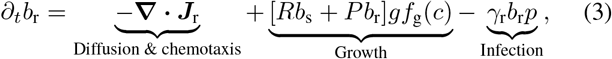

Infected less susceptible bacteria:

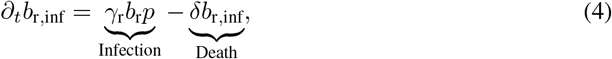

Phage:

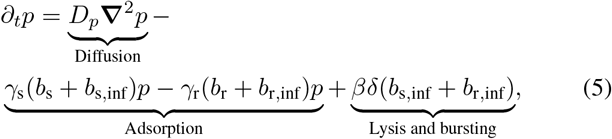

Nutrient:

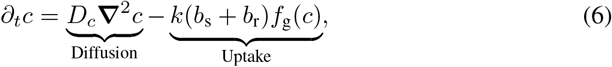

Chemoattractant:

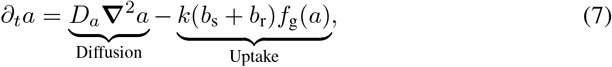

where we have neglected the contribution of chemoattractant uptake to growth since *c* » *a*.

### Model recapitulates experimental observations of front formation and steady-state dynamics

We solve Eqs. (1)-(7) numerically, with no-flux boundary conditions for all fields and initial conditions matching the experiments [Table **1**, figures S6 to S9]. As shown in Figs. **3**b-c and Videos S2-S4, the simulations, despite their simplicity, recapitulate all the key features of the experimental observations: (i) delays in chemotactic front formation that increase with phage concentration, (ii) steady-state front speeds and widths that remain unchanged despite phage predation, and (iii) sustained bacterial concentration within migrating fronts across all phage concentrations.

**Table 1.** Estimates and measurements of the parameter values of the model.

| Physical parameters | Definition | Range of values |  |
| --- | --- | --- | --- |
| $\phi$ | Void hydrogel volume fraction | 0.17 | Ref. [36] |
| $b_{s,\text{init}}$ | Initial susceptible bacterial density | $7 \times 10^{10}$ cells mL<sup>-1</sup> | Measured |
| $b_{r,\text{init}}$ | Initial less susceptible bacterial density | 1% $b_{s,\text{init}}$ | Measured |
| $p_{\text{init}}$ | Initial phage density | $10^8 - 10^{10}$ phage mL<sup>-1</sup> | Measured |
| $c_{\text{init}}$ | Initial carbon source concentration | 25 mM | Refs. [29, 36] and measured |
| $a_{\text{init}}$ | Initial chemoattractant concentration | 10 mM | Refs. [29, 36] and measured |
| $x_{\text{init}}$ | Initial thickness of bacterial inoculum | 1 mm | Measured |
| $g$ | Maximal bacterial growth rate | $0.9 \text{ h}^{-1}$ | Measured |
| $k_c = k_s = k$ | Bacterial maximal uptake rate (nutrient and chemoattractant) | $7.8 \times 10^{-11} \mu\text{mol (min cell)}^{-1}$ | Ref. [29] |
| $c_M$ | Michaelis-Menten half-velocity constant (nutrient and chemoattractant) | 3 $\mu\text{M}$ | Refs. [29, 35, 36] |
| $(a_-, a_+)$ | Characteristic sensing concentrations | (8,200) $\mu\text{M}$ | Refs. [35, 36, 49] |
| $D_b$ | Bacteria diffusivity | $1.2 \mu\text{m}^2 \text{ s}^{-1}$ | Measured |
| $D_c = D_a = D$ | Nutrient and chemoattractant diffusivity | $800 \mu\text{m}^2 \text{ s}^{-1}$ | Refs. [29, 35, 36] |
| $D_p$ | Phage diffusivity | $11 \mu\text{m}^2 \text{ s}^{-1}$ | Measured |
| $\chi$ | Chemotactic strength | $16 \mu\text{m}^2 \text{ s}^{-1}$ | Ref. [36] and fitted |
| $\gamma_s$ | Adsorption rate (susceptible) | $3 \times 10^{-12} \text{ mL phage}^{-1} \text{ min}^{-1}$ | Ref. and fitted |
| $\gamma_r$ | Adsorption rate (less susceptible) | $5 \times 10^{-3} - 10^{-2} \gamma_s$ | Fitted |
| $\beta$ | Burst size | 90 | Refs. [51–53] and fitted |
| $\delta$ | Bacterial death rate | $1 \text{ h}^{-1}$ | Refs. [53–55] |
| $R$ | Susceptible → less susceptible rate | $10^{-3}$ | Ref. [22] |
| $P$ | Growth penalty | 0.65 – 1 | Ref. [22] and fitted |
| Characteristic scales |  |  |  |
| $\ell_c = \sqrt{Da_{\text{init}}/(kb_{s,\text{init}})}$ | Characteristic length scale | 296.5 $\mu\text{m}$ | |
| $t_c = \ell_c^2/\chi$ | Characteristic time scale | 1.53 h | |
| $b_c = b_{s,\text{init}}$ | Characteristic bacterial density (all bacterial fields) | $7 \times 10^{10}$ cells mL<sup>-1</sup> | |
| $p_c = (\gamma_s t_c)^{-1} = \chi/(\gamma_s \ell_c^2)$ | Characteristic phage density scale | $1.1 \times 10^{10}$ phage mL<sup>-1</sup> | |
| $c_c = c_{\text{init}}$ | Characteristic carbon source concentration scale | 25 mM | |
| $a_c = a_{\text{init}}$ | Characteristic chemoattractant concentration scale | 10 mM | |
| Dimensionless parameters |  |  |  |
| $\tilde{\chi} \equiv (\chi/\ell_c^2)/(b_{s,\text{init}}\gamma_s)$ | Escape parameter | 0.05 | |
| $\beta$ | Burst size | 90 | |
| $\tilde{\delta} \equiv \delta t_c$ | Bacterial death rate/Bacterial chemotaxis rate | 3.06 | |
| $\tilde{g} \equiv g t_c = g/(\chi/\ell_c^2)$ | Bacterial growth rate/Bacterial chemotaxis rate | 0.9 | |
| $\text{MOI} \equiv p_{\text{init}}/(b_{s,\text{init}})$ | Multiplicity of infection | $10^{-3} - 10^{-1}$ | |
| $\gamma_r/\gamma_s$ | Less-susceptible-to-susceptible adsorption ratio | $5 \times 10^{-3} - 10^{-2}$ | |
| $b_{r,\text{init}}/b_{s,\text{init}}$ | Less-susceptible-to-susceptible initial density | $10^{-2}$ | |
| $R$ | Susceptible → less susceptible rate | $10^{-3}$ | |
| $P$ | Growth penalty | 0.65 – 1 | |
| $a_{\text{init}}/c_{\text{init}}$ | Ratio of initial concentrations | 0.5 | |
| $\tilde{D}_b \equiv D_b/\chi$ | Bacterial diffusivity/Bacterial chemotaxis | 0.075 | |
| $\tilde{D}_p \equiv D_p/\chi$ | Phage diffusivity/Bacterial chemotaxis | 0.69 | |
| $\Gamma_c \equiv \chi/D$ | Bacterial chemotaxis/Nutrient diffusivity | 0.02 | |
| $\Gamma_s \equiv \Gamma_c \equiv \Gamma$ | Bacterial chemotaxis/Chemoattractant diffusivity | 0.02 | |
| $\tilde{c}_{M,c} \equiv c_M/c_{\text{init}}$ | Dissociation constant/Initial concentration | $1.5 \times 10^{-4}$ | |
| $\tilde{c}_{M,s} \equiv c_M/a_{\text{init}}$ | Dissociation constant/Initial concentration | $3 \times 10^{-4}$ | |
| $\tilde{x}_{\text{init}} \equiv x_{\text{init}}/\ell_c$ | Dimensionless thickness of bacterial inoculum | 3.37 | |

At the lowest phage concentration (*p*_init_ = 10^8^ PFU*/*mL, MOI_init_ ∼ 10^−2^), the bacterial population has sufficient time to consume nutrients, proliferate, and establish a strong chemoattractant gradient before phages can diffuse into the bacterial population and begin killing cells [Video S2, Fig. **3**c]. The resulting chemotactic migration is fast enough to prevent the spread of infection from newly produced phage, which are left behind the advancing front [pink curve in Fig. **3**c, rightmost panel]. Consequently, the steady-state front speed, width, and bacterial concentration remain unchanged despite phage predation [Fig. **3**b-c], in good agreement with the experimental observations.

As phage concentration increases to *p*_init_ = 10^9^ and 10^10^ PFU*/*mL (MOI_init_ ∼ 10^−1^ and 10^0^, respectively), phage predation initially outpaces bacterial chemotaxis, leading to collapse of the fully susceptible subpopulation [Video S3-S4, Fig. **3**c]. The less susceptible bacteria must then proliferate to a high concentration and reestablish a chemoattractant gradient before the front can migrate, causing both the time and position of front formation to increase with phage concentration [Fig. **3**b]. However, once fronts form, they propagate at the same steady-state speed as the phage-free control across all conditions tested, in good agreement with our experimental findings.

These results demonstrate that our minimal biophysical model quantitatively recapitulates the key experimental observations and unveils the underlying mechanism: bacteria outrun the spreading infection by migrating collectively faster than phage bursts can propagate. Collective migration through a spatially-extended environment thus enables population survival, in stark contrast to well-mixed populations where the same phage concentrations rapidly eliminate susceptible bacteria.

### Dimensionless escape parameter predicts front formation dynamics

Our simulations show that whether bacteria can outrun phage depends on the competition between chemotactic migration and phage infection. To quantify this competition, we compare their characteristic rates. Bacterial chemotaxis occurs at rate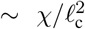, where *χ* is the chemotactic coefficient and 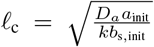 is the characteristic distance over which chemoattractant is depleted by the migrating front, which sets the front width. Phage infection depletes the bacterial population at rate ∼ *γ*_s_*b*_s,init_, determined jointly by the efficacy of phage adsorption and the density of bacteria. The ratio of these two rates defines a dimensionless *escape parameter*, 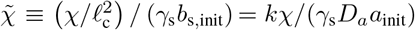. When 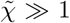, we expect that bacteria can escape phage infection using chemotaxis; conversely when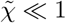, infection outpaces chemotaxis and front formation is increasingly delayed. Whether bacteria ultimately survive also depends on the initial effective multiplicity of infection MOI_init_ ≡ *p*_init_*/b*_init_; when MOI_init_ is large, we expect that phage rapidly decimate the susceptible population before a chemotactic gradient can be established, while when MOI_init_ is small, bacteria have sufficient time to proliferate and form a migrating front before widespread infection occurs. While other parameters, such as the phage burst size *β* [Fig. S10], can also influence front formation dynamics, we focus on 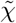 and MOI_init_ because these two dimensionless parameters are directly controllable in our experiments and capture the essential processes underlying chemotactic front formation. To determine how 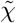 and the initial multiplicity of infection MOI_init_ jointly dictate population behavior, we perform 58 different simulations across a wide range of parameter values [Table **1**].

The resulting state diagram [Fig. **4**] reveals three distinct regimes. At high 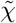 and low MOI_init_ (faster chemotaxis, fewer phage), fronts form immediately without delay [light grey, Fig. **4**], indistinguishable from the phage-free control— bacterial chemotaxis is so rapid that phage predation has negligible impact. At intermediate conditions, fronts exhibit delays in formation [medium grey] that increase with MOI_init_, as the less susceptible subpopulation must proliferate and reestablish the chemotactic gradient after the susceptible population collapses; however, once formed, these fronts propagate at the same steady-state speed as the control case. Our three experimental conditions (*p*_init_ = 10^8^, 10^9^, and 10^10^ PFU/mL at *χ* = 16 μm^2^/s) map directly onto these regimes, as shown by the pink circles in Fig. **4**. Ultimately, at sufficiently low 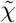 and high MOI_init_ (slower chemotaxis, abundant phage), we expect that phage predation outpaces chemotaxis, preventing front formation entirely—the front formation time diverges and the migrating population collapses.

**Fig. 4.**
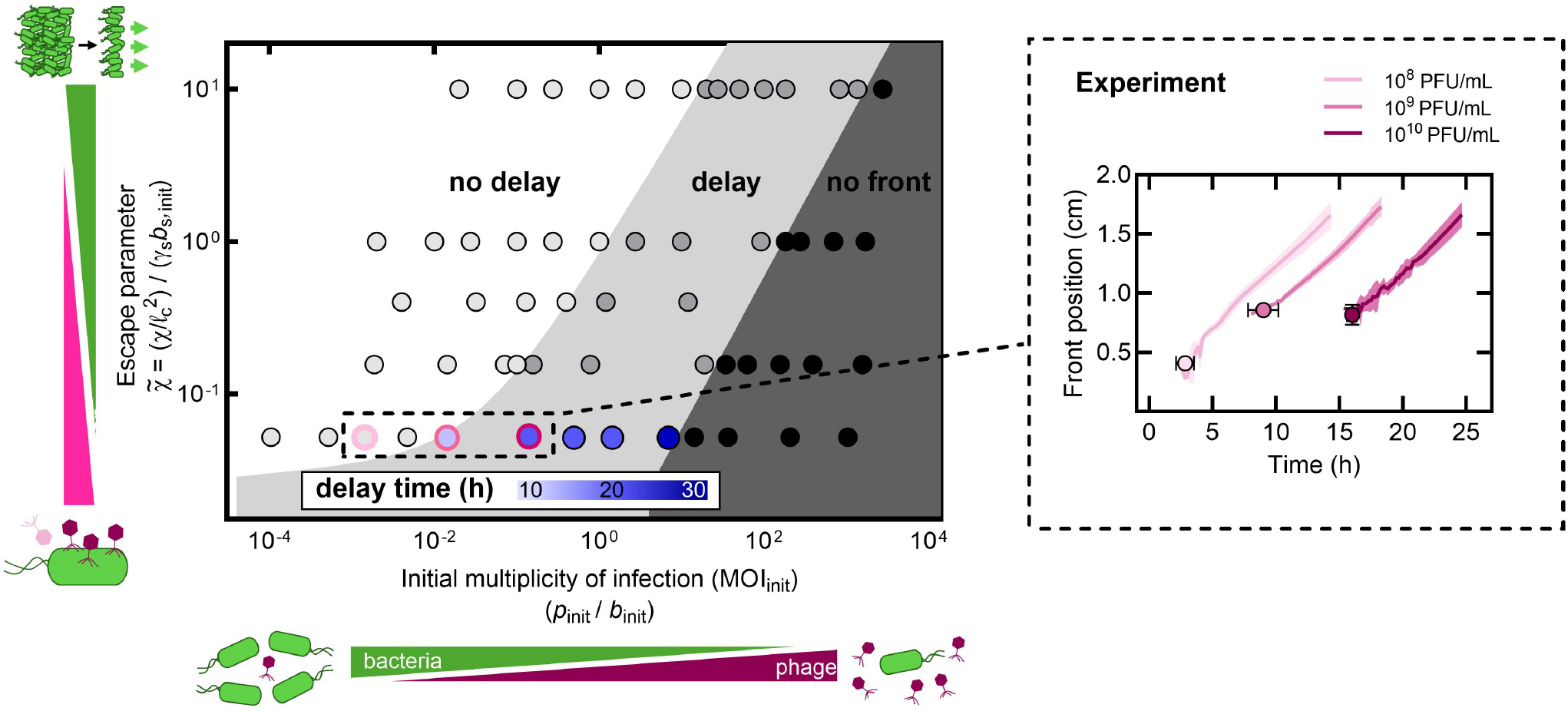
State diagram shows how the escape parameter 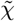 and initial multiplicity of infection MOI_init_ influence chemotactic front formation timing. State diagram from biophysical model simulations showing how chemotaxis, phage predation, and initial phage-to-bacteria ratio affect front formation. Vertical axis: dimensionless escape parameter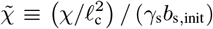, which compares the rates of chemotaxis and phage infection; ℓ_c_ is the distance over which chemoattractant penetrates into the migrating front. Horizontal axis: initial multiplicity of infection MOI_init_ = *p*_init_*/b*_s,init_. As MOI_init_ increases, phage predation increasingly delays front formation, eventually preventing formation entirely. As 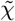 increases, bacteria increasingly outrun phage via enhanced chemotactic efficiency, reducing delays. While other parameters of the model, such as the phage burst size or bacterial lysis time, can affect the position of the boundaries delimiting these regions, we expect the same qualitative behavior. The boundary delimiting the no delay-delay regions is well described by the following empirical function: 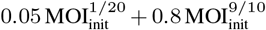, whereas the boundary delimiting the delay-no front regions is approximately described by the empirical scaling 0.006 MOI_init_. For a fixed value of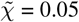, the delay time predicted by model simulations is shown as the blue-shaded regions. Light pink circle: experimental condition at 10^8^ PFU/mL showing minimal delay. Medium and dark pink circles: conditions at 10^9^ and 10^10^ PFU/mL showing increasing delays. Inset: front position over time for three experimental datasets highlighted in phase diagram and shown in Figs. **1**d and **3**b. Data represent mean ± standard deviation (*n* ≥ 3).

The hydrogel platform provides a straightforward way to further test this framework: reducing the packing density of hydrogel particles increases the sizes of the pores between them, increasing chemotactic efficiency purely physically by reducing cellular confinement [27, 36, 50]. Therefore, as a final test of the model, we repeat the experiment for

*p*_init_ = 10^9^ PFU/mL, but in looser hydrogel packings, for which *χ* increases from 16 to 40 μm^2^/s [Video S5]. The theory predicts that, in this case, a chemotactic front should form without delay [Fig. **5**a], with no influence of phage predation on front formation time or steady-state speed. Our experiments directly confirm this prediction, as shown in Figs. **5**b-d, providing additional validation of our biophysical framework. Altogether, these results establish our state diagram as a quantitative framework for predicting bacterial survival: for a given MOI_init_, increasing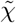, either by improving chemotactic efficiency or hindering phage infection, enables bacteria to collectively escape phage predation.

**Fig. 5.**
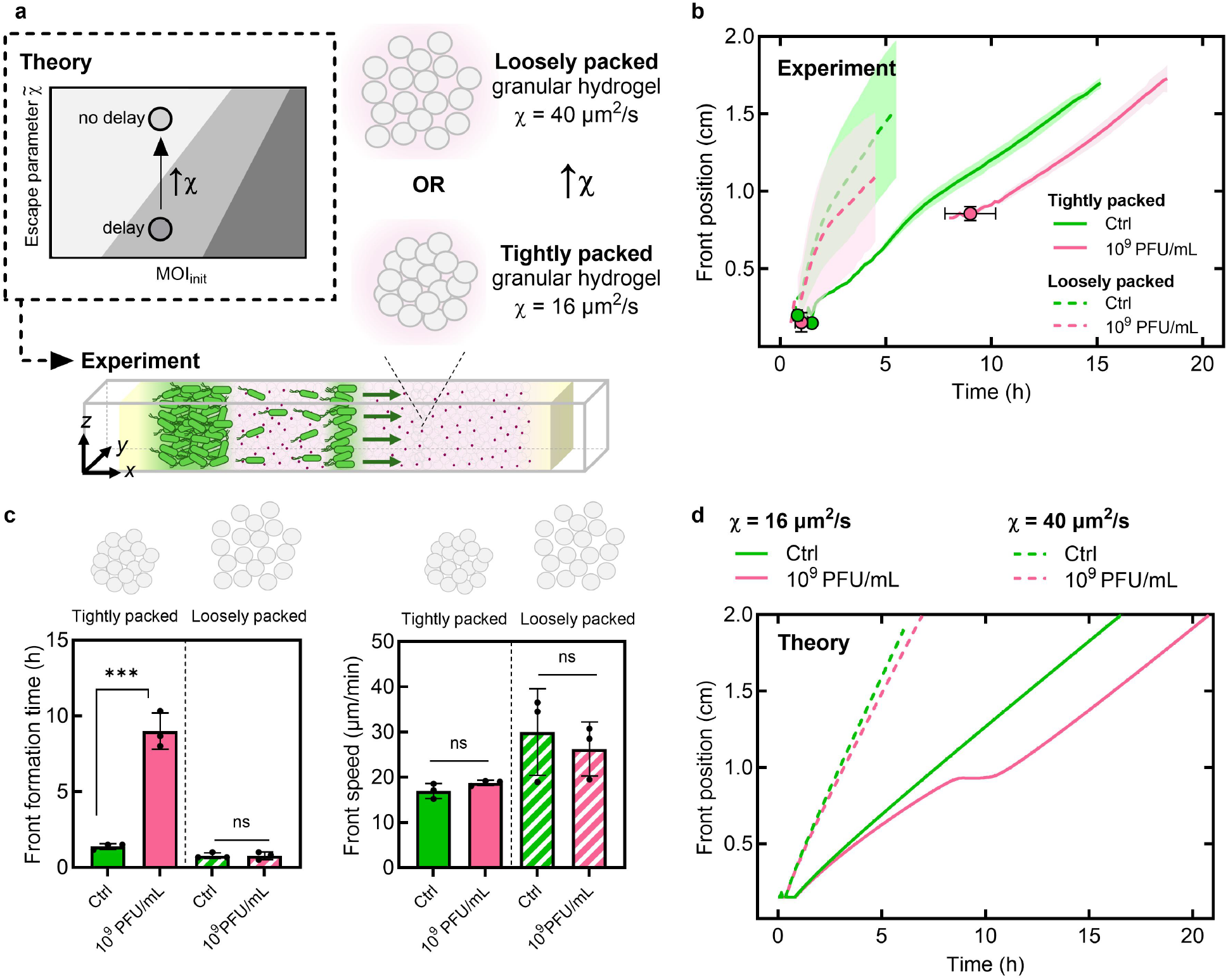
Increasing chemotactic efficiency enables escape from phage predation. **a**, Schematic showing how decreasing the hydrogel particle packing density improves bacterial chemotactic efficiency; our theory predicts that, in this case, chemotactic front formation will no longer be delayed. **b**, Bacterial front position over time in tightly versus loosely packed hydrogels, either for the no-phage control (green) or when loaded with 10^9^ PFU/mL phage (pink). **c**, Front formation time and steady-state speed without phage (control) and with 10^9^ PFU/mL phage for tightly and loosely packed hydrogels. **d**, Theoretical predictions for the same *χ* values as experiments. Model predicts no delay at *χ* = 40 *μ*m^2^/s versus 9 h delay at *χ* = 16 *μ*m^2^/s. Data represent mean ± standard deviation (*n* ≥ 3). One-way ANOVA: ns = not significant, \*\*\**p* < 0.001.

## Discussion

Our work helps to resolve the puzzle of how bacteria successfully traverse phage-rich terrain, from the human gut to ocean sediments and agricultural soils, without being decimated. We show that chemotaxis alone enables bacteria to collectively escape phage predation in spatially-extended environments by outrunning the spreading infection [Fig. **6**], without requiring evolution of genetic resistance. This finding challenges the prevailing focus on resistance as the primary mechanism enabling bacterial populations to survive phage predation—a paradigm established by studies in well-mixed liquid cultures, where resistance mutations consistently emerge and spread. Moreover, our biophysical model provides a framework that unifies the different effects of bacterial motility, chemoattractant consumption and sensing, phage infection, and the abundance of both phage and bacteria, yielding quantitative principles to predict and control these dynamics more broadly.

**Fig. 6.**
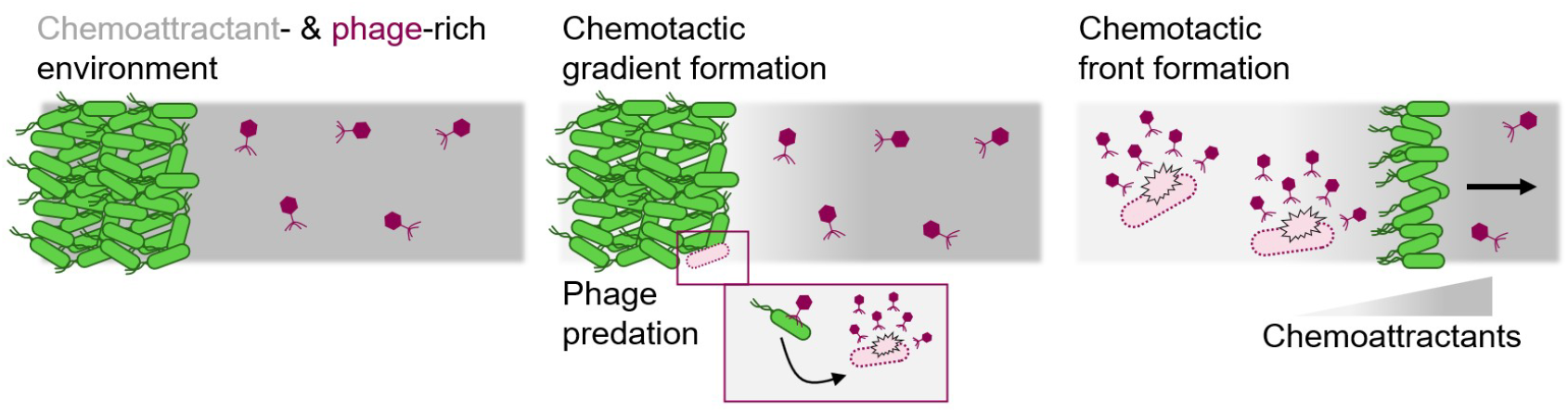
Schematic overview of findings. Bacteria inoculated at the left edge of a phage-rich environment (left) consume chemoattractant and establish a self-generated gradient (middle). Despite continuous phage infection and lysis of cells within the population, collective chemotactic migration up this gradient is fast enough to outrun the trailing burst of phage progeny, enabling the formation and propagation of a migrating chemotactic front (right). This mechanism allows bacterial populations to successfully traverse phage-rich terrain over large distances without requiring evolution of genetic resistance.

Several simplifications in the model represent opportunities for future investigation. We assumed cells transition from susceptible to less susceptible at a fixed rate, but in reality, phenotypic transitions may also depend on environmental conditions including local phage concentration, nutrient availability, and population density. Incorporating such contextdependent transitions could reveal how bacteria dynamically modulate their susceptibility in response to infection pressure. We also classified bacteria into just two discrete subpopulations, but natural populations exhibit continuous distributions of susceptibility arising from cell-to-cell heterogeneity in receptor expression and physiological state. Extending the model to include a spectrum of susceptibility states, or stochastic transitions between states, would provide a more realistic representation of population-level dynamics. Furthermore, while we described front formation dynamics using the experimentally-tunable dimensionless parameters 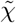 and MOI_init_, other parameters also influence population survival. For example, phage burst size *β* determines how rapidly phage populations amplify following infection [Fig. S10], while bacterial growth rate *g* affects how quickly the population can recover from infection events. Thus, alternative dimensionless groups may provide additional physical insight. Exploring such alternative parameterizations will be a useful direction for future research.

Experimentally, our findings motivate new approaches to directly assess phenotypic changes during migration. RNA sequencing or single-cell transcriptomics at defined spatial positions along chemotactic fronts could identify molecular signatures associated with transient or more permanent susceptibility changes. Time-lapse imaging of individual cells expressing fluorescent phage-receptor proteins could reveal how receptor expression varies spatially within migrating fronts and whether this variation correlates with infection probability. Finally, while our experiments used *E. coli* and T4 phage as a model system, our transparent hydrogel platform enables study of diverse phage-bacteria pairs across different ecological contexts, from pathogens in clinical settings to environmental microbes in soil and sediment.

These findings reveal a previously unappreciated ecological function of bacterial chemotaxis. Chemotaxis is traditionally understood as a foraging strategy for navigating toward nutrients and away from toxins, or for promoting range expansion of a bacterial population [12–17]. Our work identifies an additional ecological function: predator evasion through collective spatial dynamics, enabling bacterial populations to persist under intense predation pressure without the fitness costs of evolving genetic resistance. As quantified by the escape parameter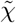, bacterial strains with higher chemotactic efficiency or lower phage susceptibility are more likely to successfully colonize distant locations, helping to inform applications in bioremediation and industrial fermentation. Conversely, in the context of phage therapy, pathogenic bacteria could escape phage via collective migration through tissues, suggesting treatment strategies that account for spatial dynamics (e.g., combining phage with motility inhibitors or engineering phage with enhanced diffusivity through capsid modifications). More broadly, we conjecture that the interplay among distinct selective pressures for nutrient acquisition, territorial expansion, and predator evasion jointly shape the evolution of motility and chemotactic sensitivity across diverse bacterial species and environmental settings.

## Acknowledgements

It is a pleasure to acknowledge insightful discussions with Grace Beggs, Lawrence Abad, Ofer Kimchi (Princeton), and members of the Datta lab. We thank Corey Stevens and Katharina Ribbeck (MIT) for assistance with experimental protocols, and the laboratory of Mark Brynildsen (Princeton) for providing the *E. coli* strain. This work was supported in part by Princeton University through the Center for the Physics of Biological Function (A.M-C and N. S.W.). This research was also supported by the National Institute of General Medical Sciences (NIGMS) of the National Institutes of Health (NIH) under Award Number R01 GM082938 (N.S.W.). The content is solely the responsibility of the authors and does not necessarily represent the official views of the National Institutes of Health. S.S.D. acknowledges support from NSF grants CBET-1941716, DMR-2011750, and EF-2124863 as well as the Camille Dreyfus Teacher-Scholar and Pew Biomedical Scholars Programs and the Princeton Catalysis Initiative. A.M.C. was supported by a Princeton Center for Theoretical Science (PCTS) fellowship and a Human Frontier Science (HFSP) fellowship (LT000035/2021). V.G.M. acknowledges funding from the Princeton Presidential Postdoctoral Research Fellow program.

## Author contributions

V.G.M., A.M-C., N.S.W., and S.S.D. designed the overall research project; V.G.M. performed all experiments and experimental analyses under the supervision of S.S.D.; A.M-C. developed the theoretical model and performed all calculations in collaboration with N.S.W. and S.S.D.; V.G.M., A.M-C., N.S.W., and S.S.D. analyzed data and wrote the paper.

## Methods

### Materials and cell culture

We use a strain of *E. coli* MG1655 that constitutively expresses GFP. T4 phage (ATCC) was propagated using standard protocols to create a 10^11^ PFU/mL stock solution in SM buffer. Reagents were from Sigma-Aldrich unless noted.

### Preparation of phage-loaded granular hydrogels

We prepare granular hydrogel packings by mixing dry granules of crosslinked acrylic-acid based co-polymers (Carbomer 980, Lubrizol, Piscataway, NJ) in liquid EZ Rich media (Teknova, Hollister, CA), as described previously [27, 29, 31, 36]. The defined medium contains *L*-serine, the primary chemoattractant, at a concentration of 10 mM. In addition, we supplement the medium with 10 mM CaCl_2_ and 10 mM MgCl_2_ following conventional protocol for enhanced phage infection efficiency. We use 10 M NaOH to adjust the final pH to 7.4 for optimal cell viability.

Using a stock solution of T4 phage in SM buffer (50 mM Tris-HCl, 8 mM MgSO_4_, 100 mM NaCl, and 0.01% gelatin) at a concentration of 1 × 10^11^ PFU/mL, we prepare phage-loaded granular hydrogel packings by reducing the stock concentration 10-, 100-, or 1000-fold by gentle mixing in granular hydrogel media, resulting in final phage concentrations of 10^8^, 10^9^, or 10^10^ PFU/mL, respectively. Phage-loaded granular hydrogel packings are then mixed with a pipette tip or inoculum wand for 1–2 minutes to ensure uniform distribution. We prepare tightly packed granular hydrogel packings at a final polymer concentration of 0.6 wt.% and loosely packed granular hydrogel packings at a final polymer concentration of 0.55 wt.%.

### Rheological characterization of granular hydrogel packings

To characterize rheological properties, we load 2 mL of granular hydrogel between roughened parallel plates (50 mm diameter) with a gap of 1 mm on an oscillatory shear rheometer (Anton Parr MCR301). We determine the storage (*G*′) and loss (*G*″) moduli under low strain (1%) at a frequency sweep of 10^−3^ – 100 Hz, and we perform a unidirectional shear rate ramp from 10^−2^ – 10^3^ s^−1^ to characterize yield stress behavior.

### Determining phage stability in granular hydrogels

To demonstrate the stability of T4 phage within granular hydrogels at 30°C, we prepare a phage-free and a phage-loaded granular hydrogel (10^9^ PFU/mL) and incubate it at 30°C for 24 h. The next day, we dilute a stationary phase culture of *E. coli* (OD ∼ 1.2) 100-fold into the pre-incubated phage-free and phage-loaded granular hydrogels. We also prepare fresh phage-free and phage-loaded granular hydrogels and dilute a stationary phase culture of *E. coli* 100-fold into freshly prepared granular hydrogels as well. Using a pipette tip, we gently mix *E. coli* into granular hydrogels to ensure uniform distribution. Using a plate reader (Agilent BioTek), we measure the optical density at 600 nm in a 96-well plate every 10 minutes for 24 h while incubating at 30°C. We compare bacterial growth in freshly prepared and pre-incubated phage-free granular hydrogels, as well as killing dynamics in freshly prepared and pre-incubated phage-loaded granular hydrogels. Pre-incubation at 30°C does not impact *E. coli* growth in phage-free granular hydrogels, nor does it impact phage-killing dynamics in phage-loaded granular hydrogels, demonstrating the stability of T4 phage in our granular hydrogel system at 30°C for at least 48 h.

### Setting up chemotaxis assays in glass capillaries

To set up chemotaxis assays, we use square borosilicate glass capillaries (VitroCom) with an inner diameter of 3 mm cut to a length of 3 cm. We transfer ∼170 *μ*L of phage-loaded granular hydrogel into the center of the capillary using a pipette. We prepare a dense suspension of *E. coli* by centrifuging 2 mL of an exponential phase culture (OD ∼ 0.6) made in 2 wt.% LB media at 3000 rpm for 8 min, then subsequently removing the supernatant and resuspending the bacterial pellet in 100 *μ*L of EZ Rich media, resulting in a dense suspension with a concentration of ∼ 1 × 10^10^ CFU/mL. We transfer 10 *μ*L of the dense cell suspension to the left-hand side of the glass capillary, creating a defined bacteria–granular hydrogel interface. We seal both sides of the glass capillary with ∼30 *μ*L of oxygen-permeable paraffin oil to prevent evaporation. Filled glass capillaries are secured onto long microscope cover glass (74 mm x 32 mm) with parafilm and transferred to the confocal microscope for imaging.

### Confocal microscopy chemotaxis assays

To image bacterial migration in phage-loaded granular hydrogels, we use a Nikon A1R inverted laser-scanning confocal microscope with a stage-top warming plate set to 30°C. Over the length of the capillary, we take three vertical planar images with a step size of 100 *μ*m. We take images every 20 min for up to 30 h. For post-processing, we take maximum intensity z projections and tile images across the length of the capillary for use in subsequent analysis.

### Analyzing chemotaxis

We use custom MATLAB scripts to track the position, width, and intensity of the chemotactic fronts over time. We do this by averaging GFP signal intensity in the direction perpendicular to chemotactic front migration across the length of the capillary, where the chemotactic front appears as a local peak in signal. We track peak location to map front position, peak width at half max to track front width, and normalized maximum peak signal to track front intensity over time.

### Bacterial growth and phage predation in well-mixed granular hydrogels

To assess *E. coli* growth in well-mixed environments, we dilute a stationary phase culture of *E. coli* (OD ∼ 1.2) prepared in 2 wt.% LB by 100-fold into phage-free and phage-loaded granular hydrogels (10^8^ – 10^10^ PFU/mL). We gently mix with a pipette to ensure uniform distribution. Using a plate reader (Agilent BioTek), we measure the optical density at 600 nm in a 96-well plate every 10 minutes for 24 h while incubating at 30°C.

### Assaying for phage resistance in chemotactic fronts

To assess the possibility of phage resistance in chemotactic fronts, we prepare chemotactic front assays in glass capillaries and incubate at 30°C until chemotactic fronts have moved a distance of ∼1 cm. This occurs after ∼7 h for phage-free and 10^8^ PFU/mL T4 concentrations, ∼11 h for 10^9^ PFU/mL T4 concentrations, and ∼19 h for 10^10^ PFU/mL T4 phage concentrations. Chemotactic waves are visible in the capillaries by eye. To prepare cell stocks of bacteria isolated from chemotactic fronts, we use a pipette to remove 5 *μ*L of granular hydrogel at the location of the chemotactic front, dilute into 16% glycerol stocks, and freeze at −80°C for future use.

### Determining phage diffusion coefficients in granular hydrogels

To determine the diffusion coefficient of T4 phage in our granular hydrogels, we fluorescently label T4 phage with SYBR Gold stain (Fisher Scientific) by mixing 1 mL of phage stock solution with 1 *μ*L of SYBR Gold. We incubate at 4°C for 30 min in a dark environment, then subsequently add 2 mL of purification buffer (20 wt.% PEG-8000, 2 M NaCl) and incubate at 4°C overnight to crash phage out of solution.

### Statistical analysis

We report all data as mean ± standard deviation, unless otherwise indicated. We conduct statistical analysis in GraphPad Prism 9 using one-way ANOVA and a Tukey’s post hoc comparison, unless otherwise indicated. For all samples, *n* ≥ 3, \**p* < 0.05, \*\**p* < 0.01, \*\*\**p* < 0.001, \*\*\*\**p* < 0.0001, ns = not significant.

### Non-dimensionalization of the model

To reduce the number of parameters of our model, we scale all variables using the following scales for length, time, bacterial density, nutrient and chemoattractant concentration, and phage density, respectively:

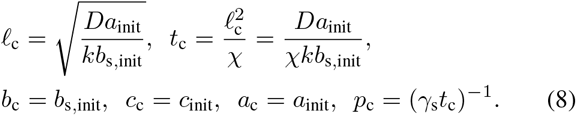

We use the initial bacterial concentration of susceptible bacteria, *b*_s,init_ to scale all bacterial density fields. The emerging dimensionless parameters as well as the values of both model parameters and dimensionless parameters used in simulations are given in Table **1**.

## SUPPLEMENTARY MATERIAL

**Video S1**. Representative confocal microscopy time lapse video of chemotactic front formation along the capillary for a control case (no phage, top) and increasingly high phage concentration (10^8^ - 10^10^ pFU/mL). Scale bar = 1 mm.

**Video S2**. Normalized model number densities of bacterial and phage populations, and chemoattractant concentration as functions of position *x* over time for an initial phage concetration of 10^8^ PFU/mL.

**Video S3**. Same as Video S2 but for initial phage concentration 10^9^ PFU/mL.

**Video S4**. Same as Video S2 but for initial phage concentration 10^10^ PFU/mL.

**Video S5**. Representative confocal microscopy time lapse video of chemotactic front formation along the capillary for a control case (no phage, top) and intermediate phage concentration (10^9^ PFU/mL) in loosely packed granular hydrogel. Scale bar = 1 mm.

**Fig. S1.**
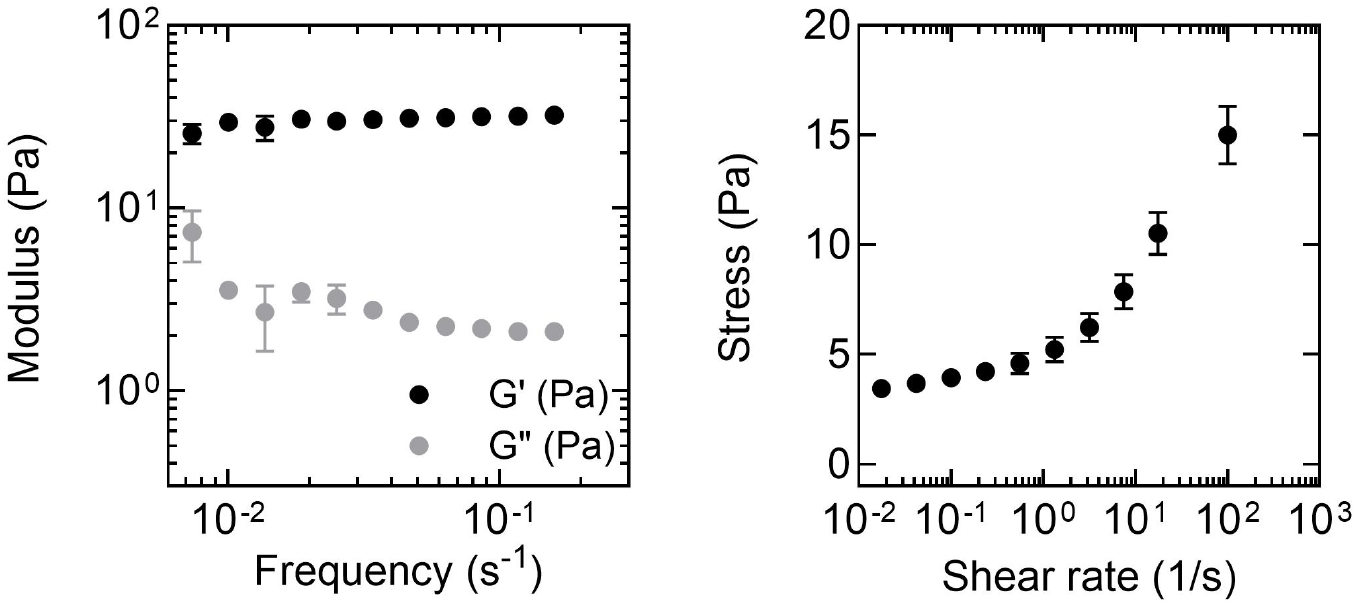
Rheological characterization of granular hydrogels (0.6 wt.% Carbopol). Left: Storage (G’) and loss (G”) moduli as a function of frequency (10^−3^ - 1 Hz). Right: Stress as a function of shear rate (10^−2^ – 10^3^ s^−1^). Data represent mean ± standard deviation (*n* ≥ 3).

**Fig. S2.**
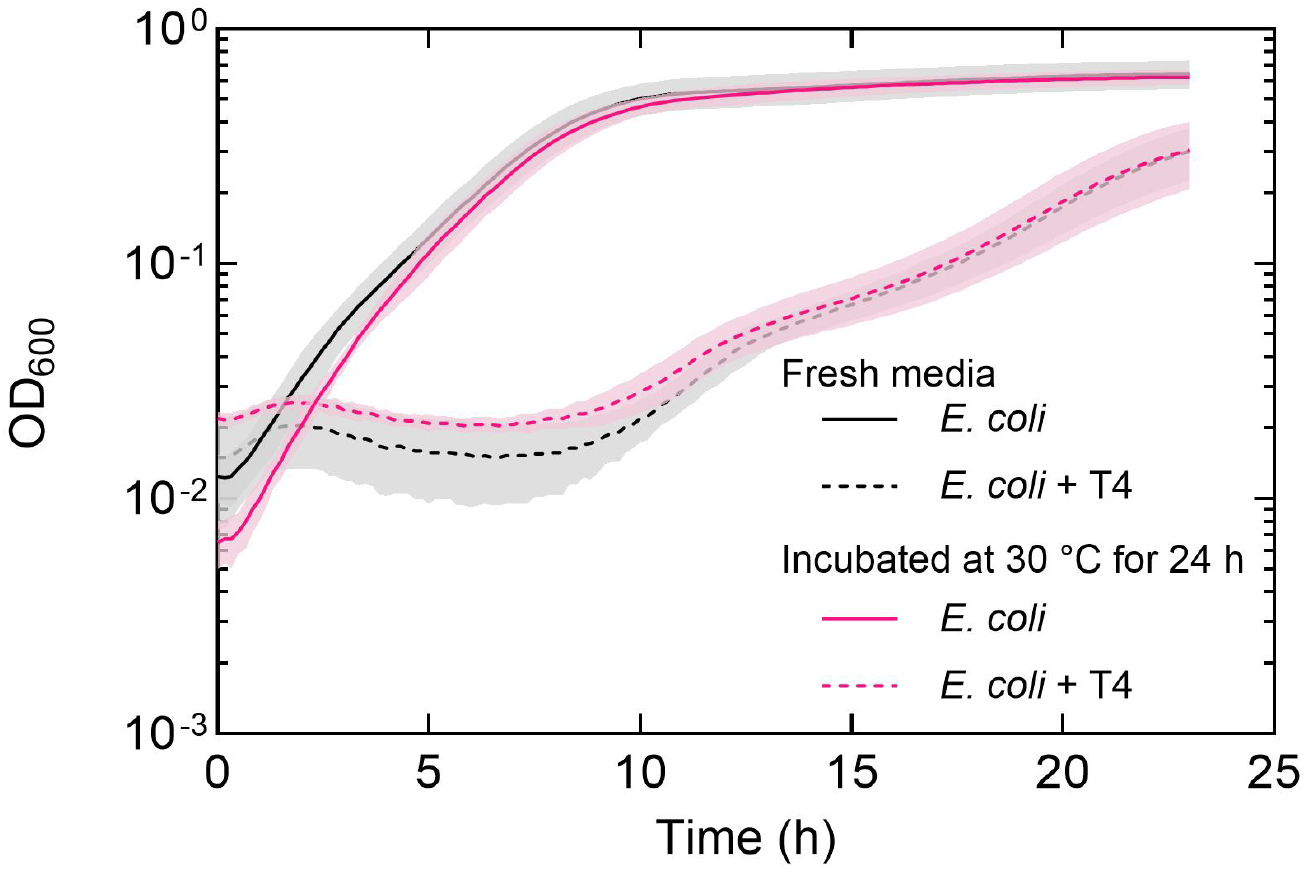
Phage stability in granular hydrogels. Optical density at 600 nm (OD600) of *E. coli* growing in granular hydrogels (0.6 wt.% Carbopol) with (dotted) or without (solid) T4 phage. Black curves show OD600 over 24 hours after inoculating fresh granular hydrogels (+/-phage) with *E. coli*. Pink curves show OD600 over 24 hours after inoculating 24-hour-old hydrogels (+/-phage) incubated at 30°C. Growth (no phage) and killing dynamics (with phage) remain unchanged after hydrogels are incubated for 24 hours. We conclude that phage remain stable in granular hydrogels for at least 24–48 hours, covering the duration of experiments in this study. Data represent mean ± standard deviation (*n* ≥ 3).

**Fig. S3.**
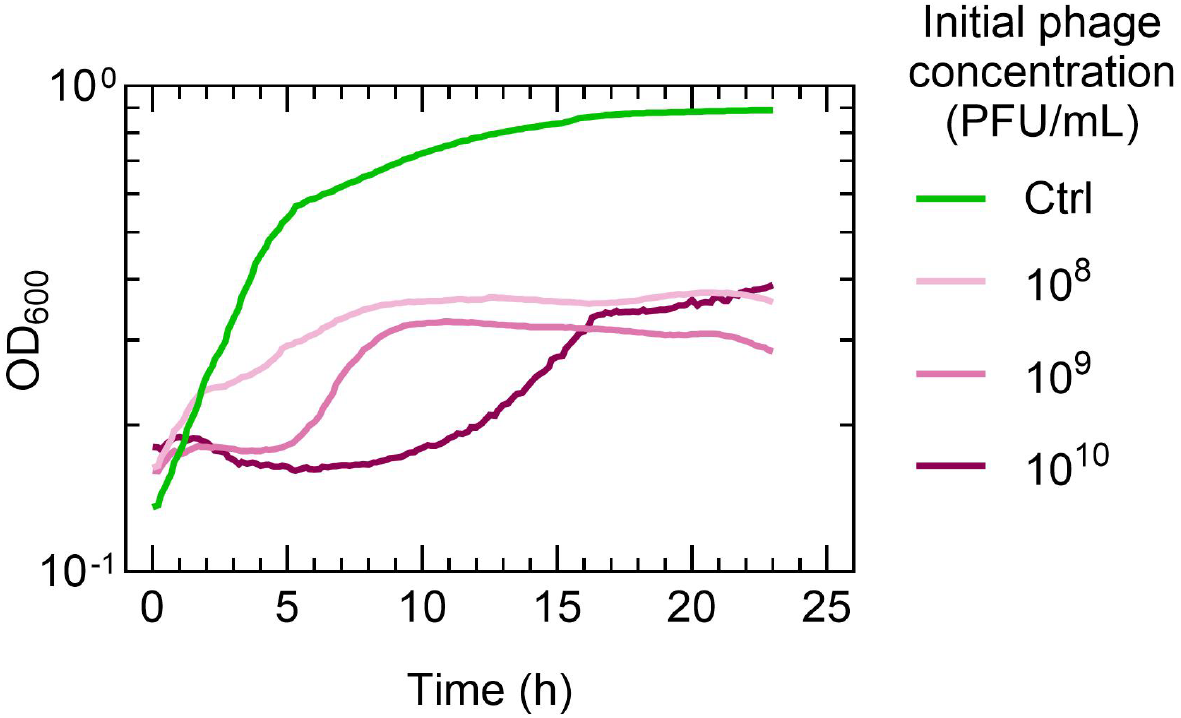
Optical density at 600 nm (OD600) of *E. coli* growing in hydrogels (0.6 wt.% Carbopol) at various phage concentrations in a well-mixed environment. *n* = 1.

**Fig. S4.**
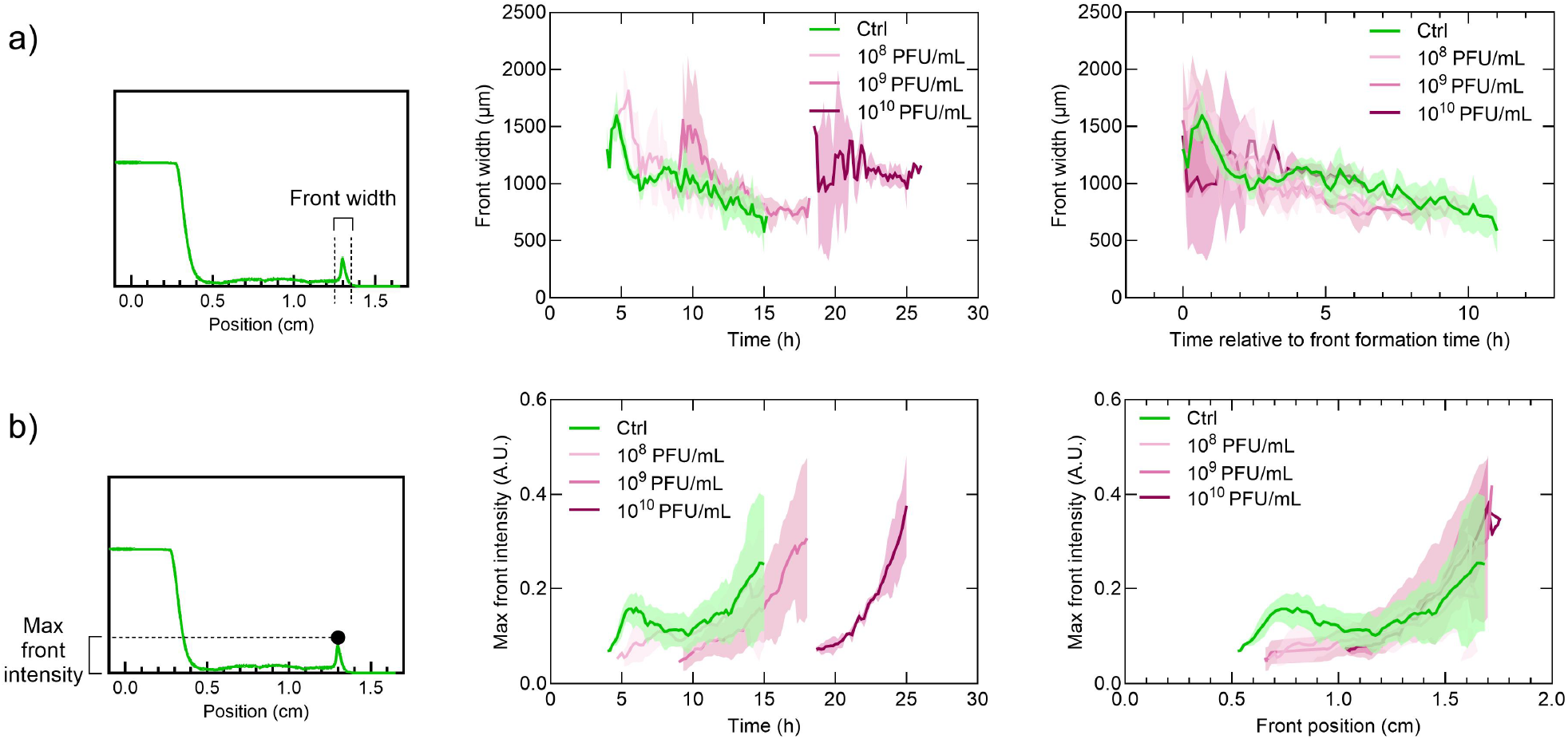
Characterizing front width and front intensity over time. (**a**) Width of chemotactic fronts over time at various phage concentrations. We plot the width as a function of time and relative to the initial front formation time. (**b**) Maximum fluorescence intensity of chemotactic fronts over time, plotted as a function of time and as a function of front position. Data represent mean ± standard deviation (*n* ≥ 3).

**Fig. S5.**
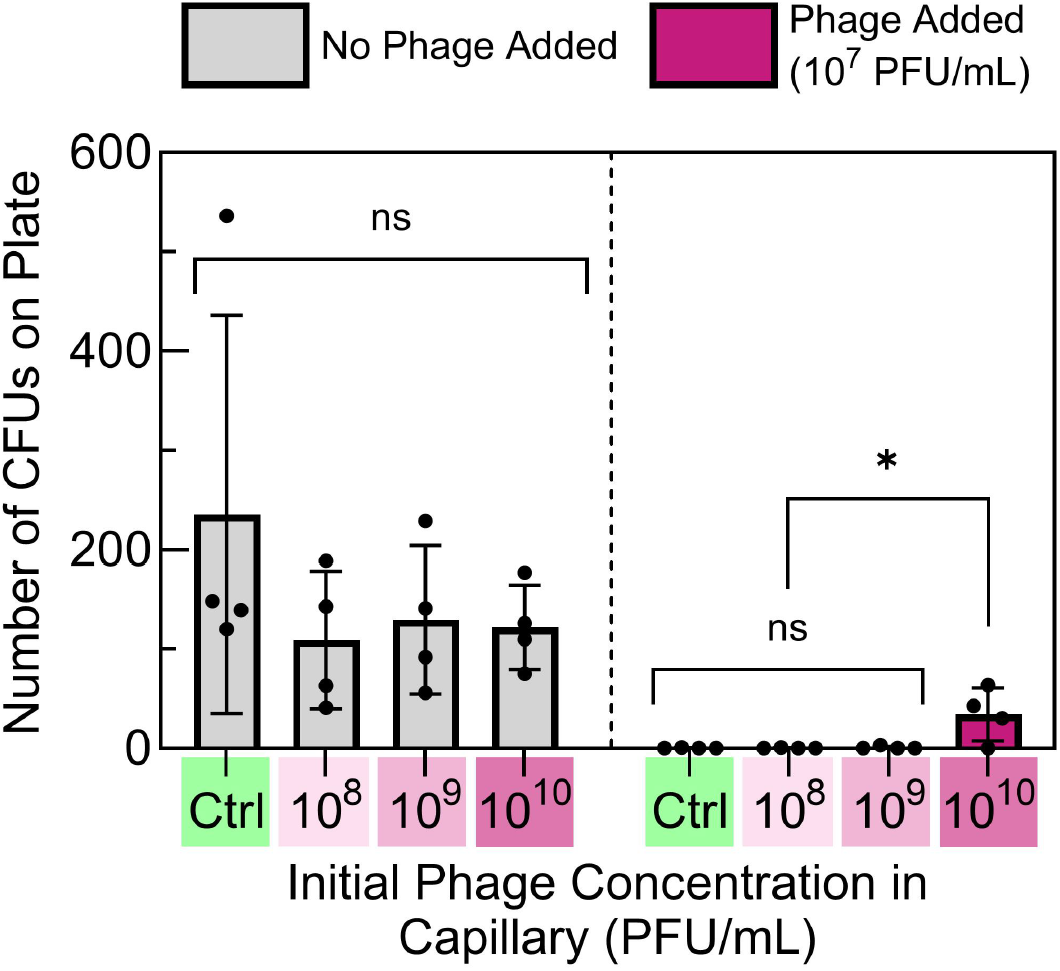
Total colony-forming units (CFUs) isolated from chemotactic fronts formed in the presence of phage. Total CFUs from chemotactic fronts plated on soft agar without phage (light gray) and with phage (dark pink, 10^7^ PFU/mL). Data represent mean ± standard deviation (*n* ≥ 4). We use one-way ANOVA to determine statistical significance: ns = not significant, \**p* < 0.05.

**Fig. S6.**
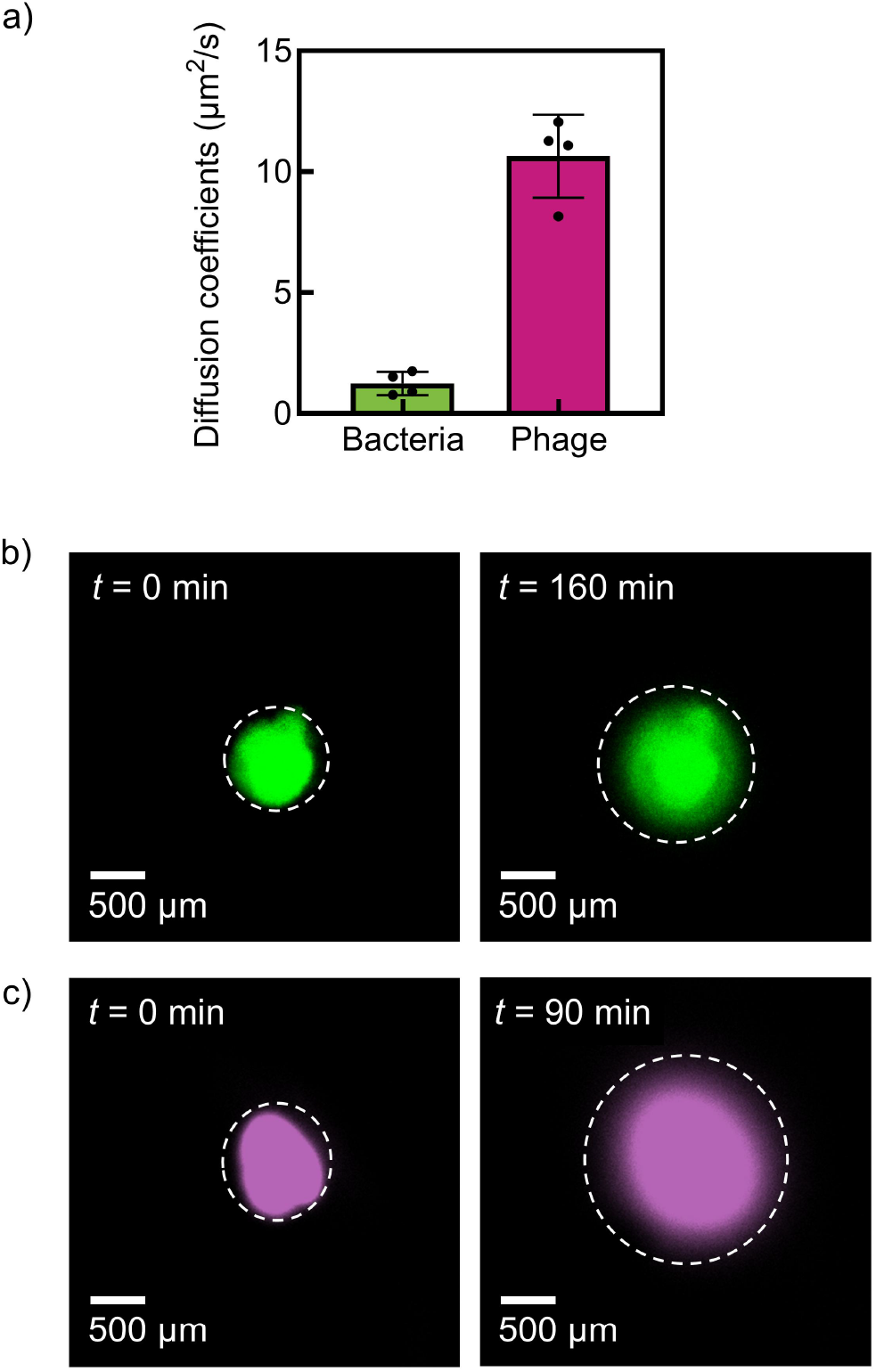
Diffusion coefficients of motile bacteria and phage in granular hydrogels (0.6 wt.% Carbopol). (**a**) Diffusion coefficients of bacteria and phage measured in granular hydrogels experimentally. (**b**) Representative maximum projection confocal *z*-stack images of bacteria diffusing in granular hydrogels at *t* = 0 min and *t* = 160 min (prior to chemotactic front formation). Scale bar = 500 *μ*m. (**c**) Representative maximum projection confocal *z*-stack images of fluorescently labeled phage diffusing in granular hydrogels at *t* = 0 min and *t* = 90 min. Scale bar = 500 *μ*m. Data represent mean ± standard deviation (*n* ≥ 4).

**Fig. S7.**
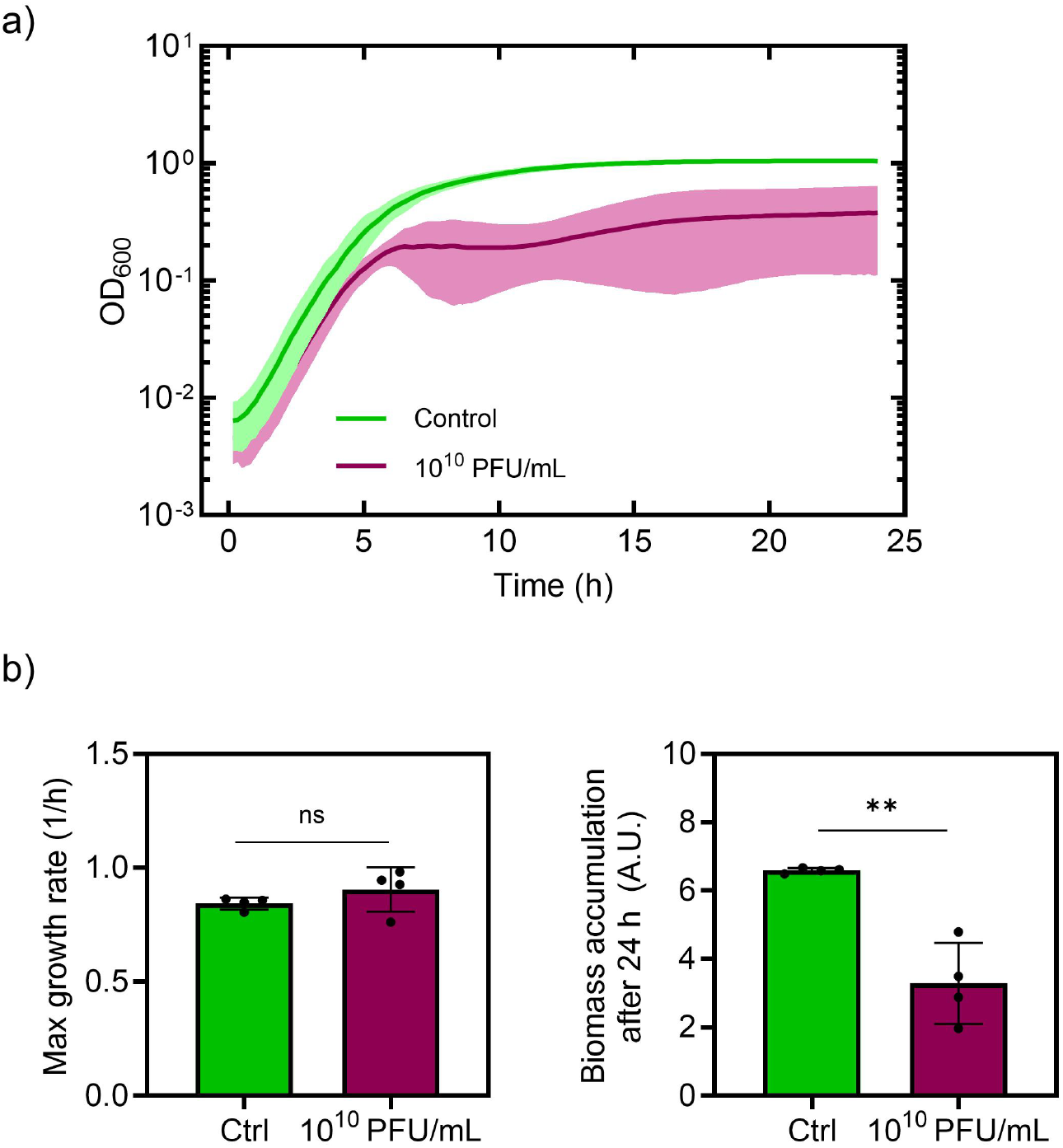
Growth reduction of *E. coli* less susceptible to phage infection. (**a**) Optical density at 600 nm (OD600) of *E. coli* isolated from chemotactic fronts formed without phage (green) and *E. coli* isolated from chemotactic fronts formed in the presence of 10^10^ PFU/mL phage (pink). (**b**) Quantification of maximum growth rate and biomass accumulation (area under the curve) after 24 h for *E. coli* isolated from chemotactic fronts. Data represent mean ± standard deviation (*n* ≥ 3). We use one-way ANOVA to determine statistical significance: ns = not significant, \*\**p* < 0.01.

**Fig. S8.**
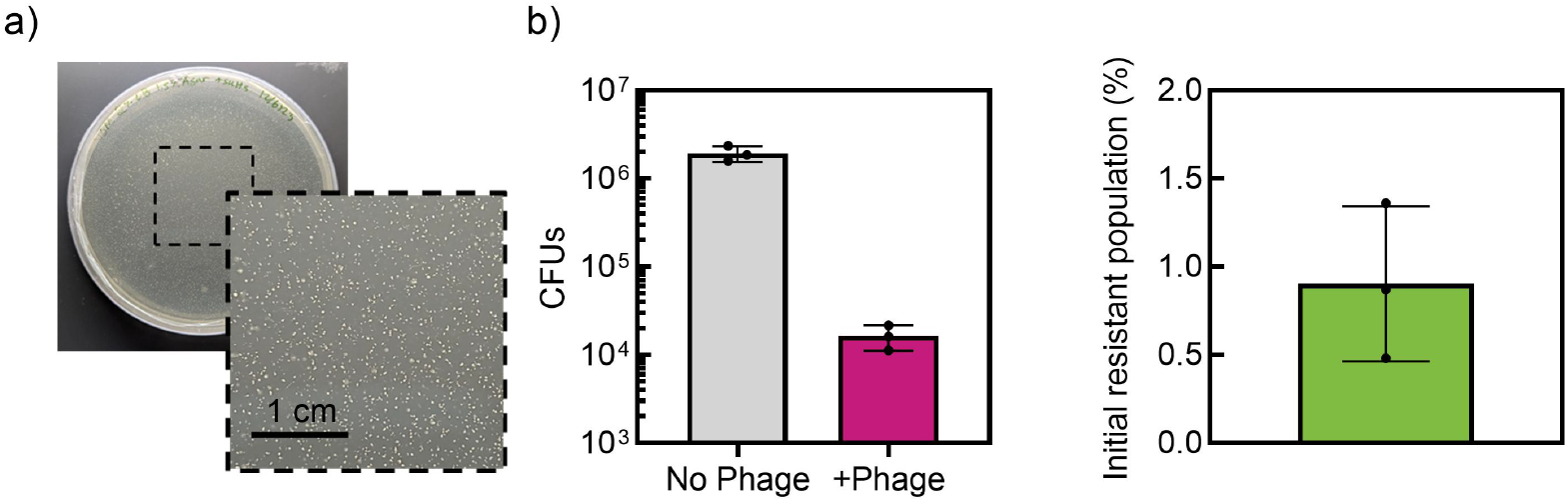
Initial fraction of phage-resistant *E. coli* in stock solutions. (**a**) Representative image of CFUs from stock solution growing on agar plates with T4 phage. (**b**) Quantification of total CFUs from stock solutions plated on phage-free (grey) and phage-rich plates (pink), and the percentage of phage-resistant bacteria in the stock population. Data represent mean ± standard deviation (*n* ≥ 3).

**Fig. S9.**
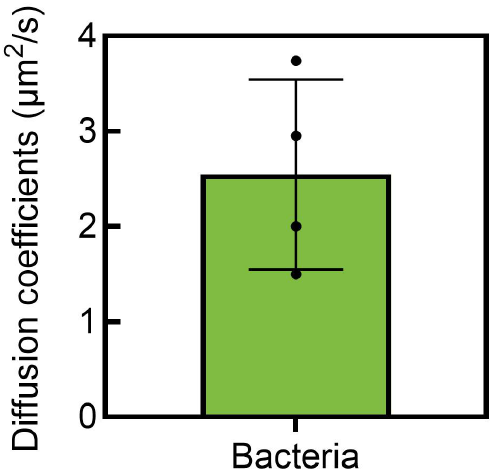
Diffusion coefficients of bacteria in measured experimentally in granular hydrogels (0.55 wt.% Carbopol). (*n* ≥ 4).

**Fig. S10.**
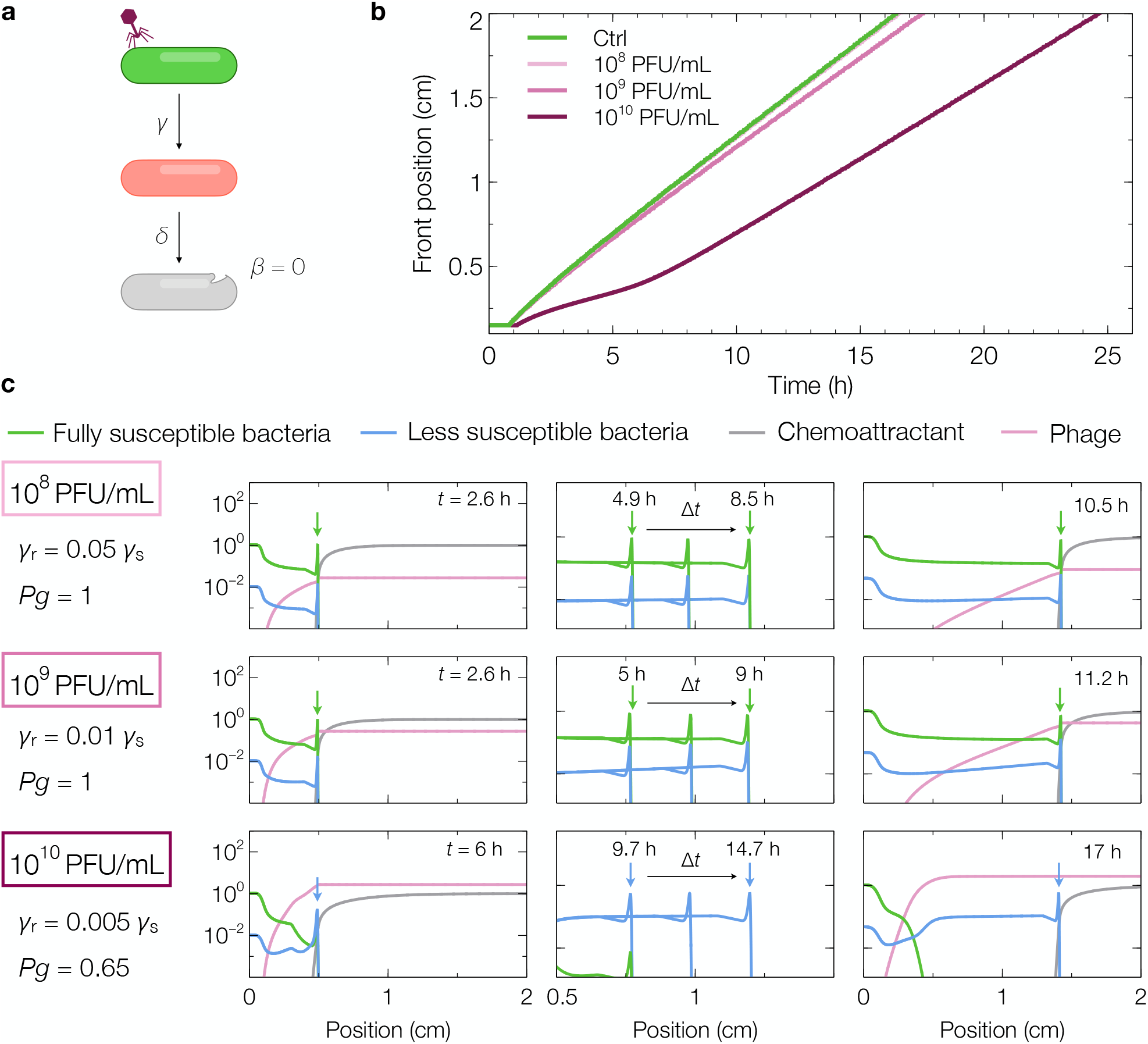
Phage burst size *β* influences bacterial chemotactic front formation dynamics. **(a)** Schematic of a modified model (relative to the one presented in the main text and Fig. **3**a) in which phages can infect and kill bacteria but do not proliferate via bursting, implying *β* = 0. **(b)** Bacterial chemotactic front position as a function of time in a phage-free environment and for *p*_init_ = 10^8^, 10^9^, and 10^10^ PFU/mL, from the model with *β* = 0, i.e., non-bursting phages. In this modified model, the chemotactic front is appreciably delayed only for *p*_init_ = 10^10^, implying that the phage burst size *β* alters the phase boundaries shown in Fig. **4. (c)** Normalized concentration profiles of bacteria, chemoattractant, and phage at different times for various phage concentrations. Bacteria and chemoattractant profiles are normalized by their initial concentrations at *t* = 0, and the phage profile is normalized by the characteristic concentration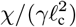, where ℓ_c_ is the chemoattractant penetration scale (see Table **1** and Methods). Initially, nutrient and chemoattractant are uniform across the domain with concentrations *c*_init_ and *s*_init_, respectively. Bacteria are uniformly distributed in a phage-free region (0 < *x* < *x*_init_) adjacent to uniform phage suspension (*x*_init_ *< x < L*), forming a slab-shaped colony facing the phage-rich environment as in the experiments. Here, *x*_init_ denotes the position of the bacteria–phage contact interface at *t* = 0, and *L* is the domain length. We assume that phage are stable and long-lived over the experimental duration and do not burst (*β* = 0), in contrast to the results shown in Fig. **3**. Arrows indicate chemotactic front position.

